# Engaging cancer patients on their attitudes towards microbiome engineering technologies

**DOI:** 10.1101/2024.11.25.625255

**Authors:** Kimberley A. Owen, Jack W. Rutter, Claire Holland, HB Trung Le, Philip Shapira, Clare Lewis, James M. Kinross, Chris P. Barnes

## Abstract

Microbiome engineering aims to develop engineered live biotherapeutic products to diagnose and treat human disease. One of the most active areas is in the engineering of modified bacteria that act as cancer therapeutics, with many products currently in clinical trials. With the emergence of any novel technologies, it is important to consult the public and end users throughout their development to address any concerns that may arise from their use. This is particularly true for therapeutics, for which it is vital that both clinicians and patients have confidence in the treatments that are available to them. Here, we surveyed a cohort of cancer patients and their relatives to gauge their knowledge of modern microbiome engineering techniques. We focus predominately on the use of bacterial eLBPs to treat cancer. Overall, most participants indicated that they would be comfortable taking cancer treatment options based on microbiome engineering technologies.

## 1 Introduction

The human microbiome is a community of microorganisms —such as bacteria, fungi, viruses, and archaea—and their genetic material that live within the body. These microorganisms coexist with their host and play critical roles in maintaining the health and functionality of that environment. Microbiome engineering aims to modify these communities in a predictable way, for example, for the treatment of diseases within the body. In principle, there are many ways to modify the microbiome, including the use of probiotics, which are food and dietary supplements containing live microorganisms. However, drugs containing live organisms that are used to treat or prevent disease are called Live Biotherapeutic Products (LBPs). Current LBPs are being used to treat a variety of conditions, including obesity, diabetes, and inflammatory bowel disease (*1*).

To improve the applicability and efficacy of LBPs, engineering biology techniques can be used to modify the microbes within the drug (*2, 3*). These are known as engineered Live Biotherapeutic Products (eLBPs). Although not yet commonly available, many eLBPs are undergoing human clinical trials, with a large fraction of these for cancer indications (*4*). In cancer treatment, eLBPs can be used for detection and reporting of biomarkers, for targeted delivery of therapeutic compounds and for eliciting an immune response (*5*). A recent review of projects from the International Genetically Engineered Machines (iGEM) competition shows the potentially diverse range of applications of eLBPs for cancer treatment (*6*). Despite this promise and the likely imminent approval of some of these products, to the best of our knowledge, there have been no previous studies that have looked at views towards engineered bacterial therapeutics or their application in cancer treatment.

As a comparison, we can look towards previous works on genetically modified (GM) foods as a proxy of general public opinion on genetically engineered products. Public attitudes towards GM foods are complex, with concerns ranging from unexpected long term health consequences to unintentional environmental harm (*7*). Furthermore, the public appear to have a greater cognizance of the potential harms of genetic modifications than they do for the benefits. In some cases, even when the benefits are known, their influence does not outweigh the negative perceptions (*8*).

Ultimately, GM foods differ from the proposed eLBPs both in composition and intended purpose. Whilst some research suggests the application of the GM product is less important than the type of gene manipulation involved (*9*), other research suggests that, within the European Union (EU), the application of GM products greatly influences people’s acceptance of the technology (*10*). It appears that the application of genetic modifications in the context of medicine meets with higher approval than other fields of genetic engineering, including GM foods (*11, 12*).

In the field of engineering biology, public acceptance has been identified as a general block to advances in the field (*13*). This stresses the need for public engagement early during research, to identify and mitigate any concerns that may not have been envisioned by the primary researchers. This is in alignment with the EPSRC Responsible Research and Innovation (RRI) initiative and recent studies by the European Commission (EC), which encouraged consulting the public on genetic engineering research (*14*).

In this study, we first perform an Early Rapid Sustainability Assessment (ERSA) of microbiome engineering as a technology (*15*). This led us to investigate public perceptions of microbiome engineering, primarily focusing on the use of engineered bacteria for cancer care. In the case of the eLBPs we consider here, the primary stakeholders will be cancer patients. The current lack of evidence exploring their views led us to launch a survey to assess attitudes towards microbiome engineering as a general principle and more specifically for the creation of novel bacterial, cancer therapeutics.

## 2 Results & Discussion

### 2.1 ERSA assessment

To understand the perception of potential future stakeholders to microbiome engineering we explored the technology through an ERSA assessment. Figure 1 shows the scores of the technology (black line). As expected, this technology scored worst under “process social”, which is the section where public acceptance is considered. The technology scores well on “Product economic” and “Process environmental” as this project is not performing worse than current standards in those areas. All remaining areas of the technology warrant further investigation.

**Figure 1.**
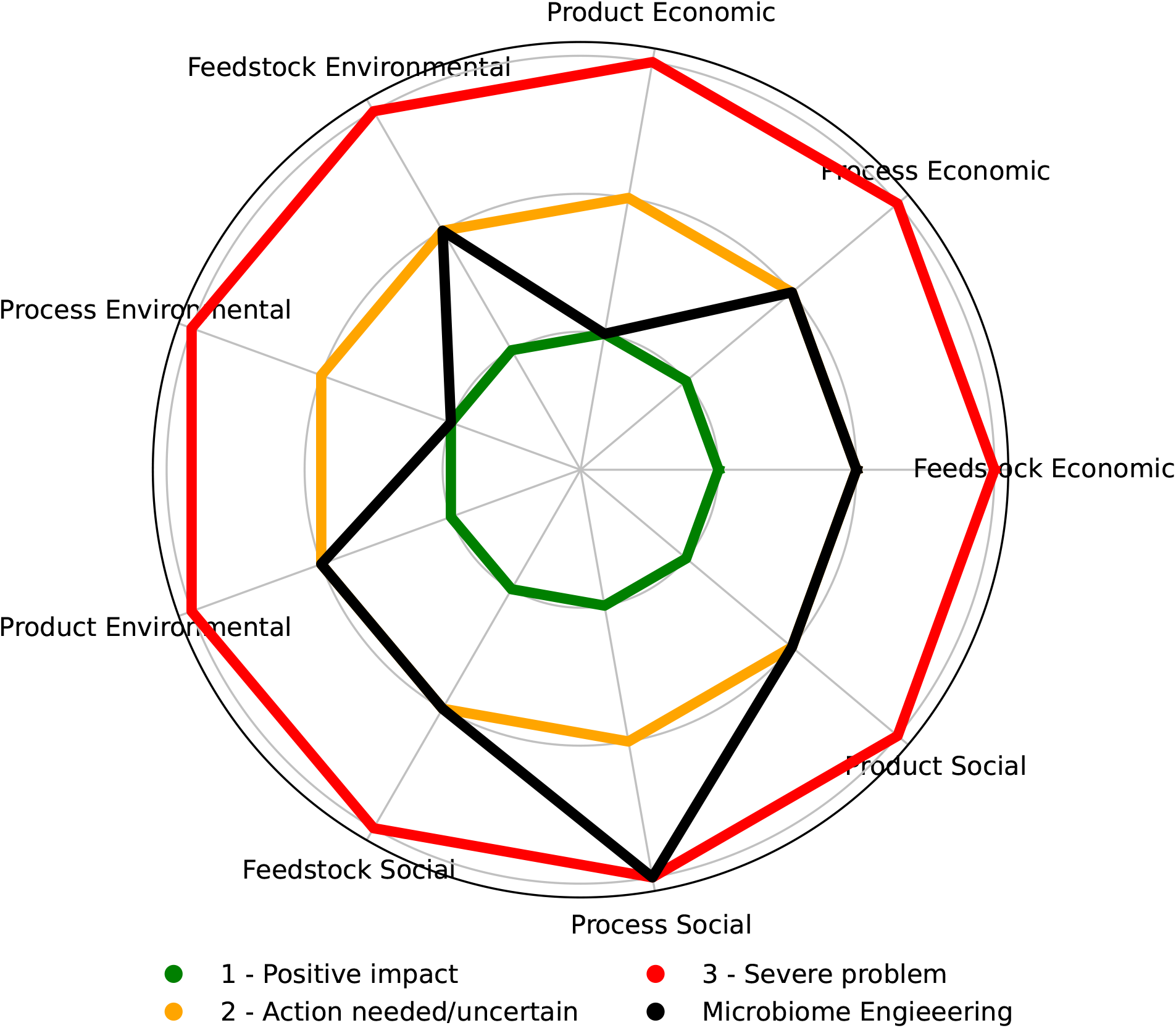
RADAR diagram constructed from the ERSA assessment of microbiome engineering. Diagram highlights areas of concern and its development. Key is given on the diagram.

### 2.2 Survey

The full survey questions are provided in the Supplementary Information. The survey received a total of 171 responses, of which 102 (59.6%) were complete. Due to the small number of responses, completed surveys from participants that had no connection to cancer (5) were removed from further analysis. More than half of the remaining participants were over the age of 60 (59%). A quarter of participants were between 40 and 59 years old (24.4%), the remaining participants were between the ages of 18 and 39 years old (16.5%). A full breakdown of these responses is given in SI Figure S1.

### 2.3 Prior knowledge of concepts

All participants were provided with definitions of the key concepts explored in the survey (these definitions are provided in the Supplementary Information) and then asked to state whether they had previously heard of the topics. The majority of respondents (95) had previously heard of probiotics. It is clear that probiotics are a well-known concept; for example the term ‘probiotic’ is often associated with dairy foods. In addition, during the initial Covid-19 outbreak, market reports predicted an increase in interest in probiotics within the food industry as the public associated probiotics with increased immune health (*16*). This was confirmed by a Chr. Hansen study, which highlighted the consumer view of probiotics and positive views on how live bacteria can benefit human health (*17*). This was further reflected by the fact that 86 respondents indicated they were aware of the proposed benefits of probiotics, wheresas only 41 were aware of the limitations. This may indicate that probiotics do not have the negative connotations that surround GM foods. A further 72 respondents reported they would be comfortable taking probiotics as part of their diet, but only 64 would be comfortable taking them to treat a medical condition (Figure 2). This conflicted with some of the open comments, which suggested greater comfortability with genetic engineering in the context of medicine. One respondent also highlighted a perception that the regulatory governance of medicine is more robust than that of food products:

**Figure 2.**
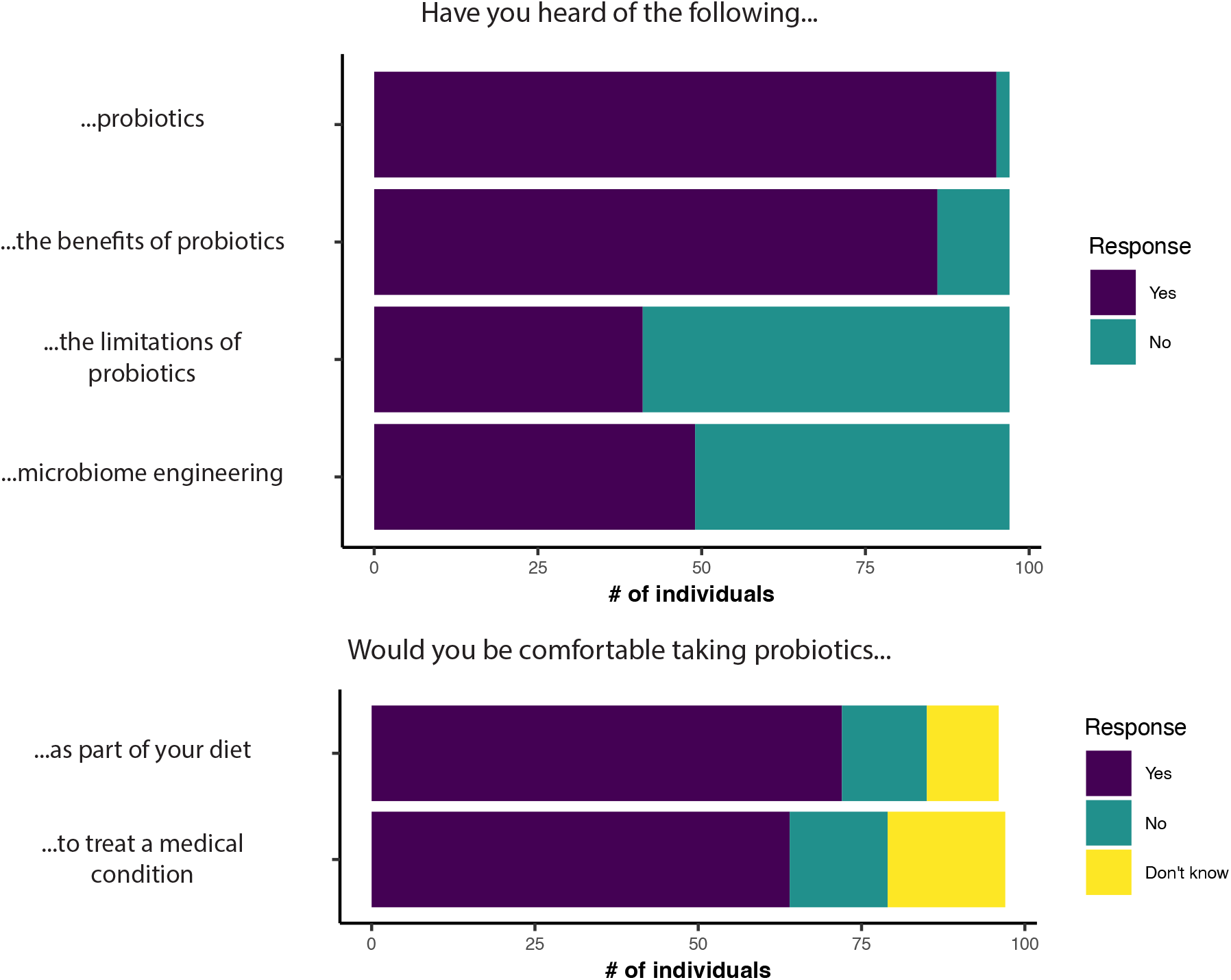
Participants where surveyed to assess their knowledge of the given topics and then asked to rank whether they would be comfortable taking probiotics as part of their diet or to treat a medical condition (n = 97 responses).

> *“I am far more comfortable as I believe the regulatory rigour / approval / testing etc to be higher”* - 2nd hand experience of cancer (40-59 age group).

In contrast to the widespread knowledge of probiotics, only 49 respondents had previously heard of microbiome engineering suggesting the term is not yet commonly known.

### 2.4 Attitudes towards current cancer therapies

Firstly, we set out to check which sources prospective patients would trust for advice on cancer treatment. Overall, doctors and clinicians were ranked as the most trustworthy source (Figure 3A). Pharmaceutical companies were ranked as the least trustworthy. This corroborates previous studies that have investigated patient trust in healthcare professionals (*18*). This is important as it highlights the sources of information patients trust and how future policymakers would be best positioned to deliver information on new therapeutics going forward. The theme of trust was prevalent in the open text answers, the word ‘trust’ was explicitly mentioned in 10 comments. Many comments focused on trust in medical teams, with respondents indicating they are open to new treatments that are recommended by their clinicians:

**Figure 3.**
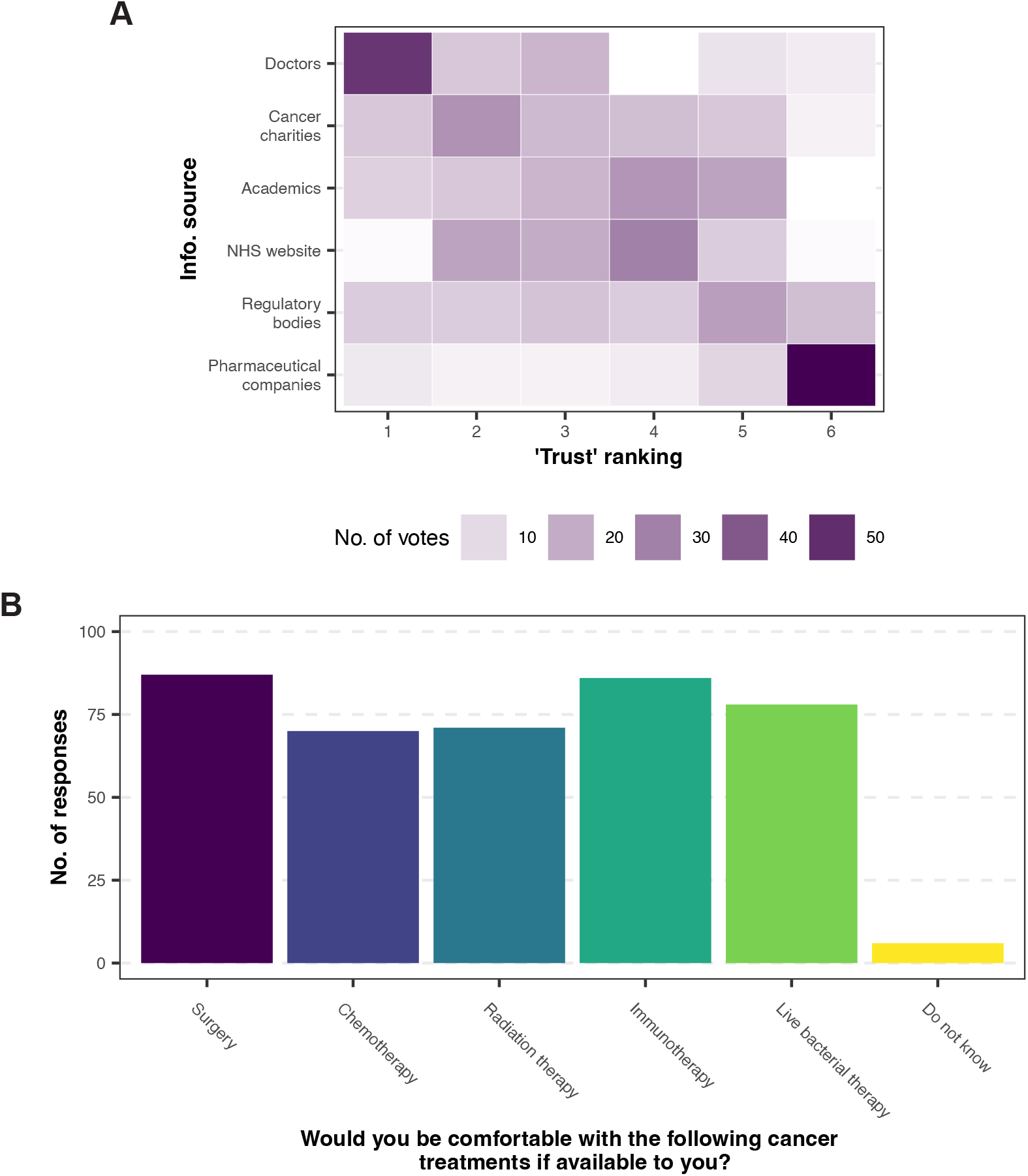
(**A**) Trust rankings for the given sources of information (a ranking of 1 was considered the most trustworthy). (**B**) How many participants would be comfortable taking the given treatment options (n = 97 responses).

> *“I’m intrigued and if my consultant recommended such treatment, I would consent to it*.*”* - 1st hand experience of cancer (40-59 age group).
>
> *“I trust the doctors, surgeon and support team (nurses, dieticians etc) to make the choice that would aim to result in the best outcome for me*.*”* - 1st hand experience of cancer (40-59 age group)
>
> *“I’d be perfectly comfortable if clinician suggested the microbiome route*.*”* - 1st hand experience of cancer (60+ age group)

Next, we explored the cancer treatment options that respondents would be comfortable taking as part of their treatment plan. Of the treatment options provided, respondents were most comfortable with surgery, closely followed by immunotherapy (Figure 3B). Chemotherapy and radiation therapy were the most unpopular options. This is likely due to awareness surrounding the adverse side effects that are associated with these treatments. Although it should be noted that over 70% of the respondents still indicated they would be comfortable with these treatments if available to them. Despite our hypothesis that live bacterial therapy would be a less popular treatment option (due to a lack of precedent for its use) a total of 77 respondents stated they would be comfortable with this treatment option. However, the issue of side effects comes up frequently in the open text comments (16 times). This links to a broader theme of social connection seen amongst the responses from participants. They are not only concerned about themselves and how they would respond to new therapies but how others would fare with new types of treatment. Particularly where the possible side effects are not yet fully known:

> *“If the microbiome can be modified in a predictable way that is beneficial to health I would be very comfortable with their use. However, most treatments have drawbacks or side effects. Without knowing what these might be I cannot be sure of whether I am comfortable with the technology*.*”* - 2nd hand experience of cancer (60+ age group).
>
> *“The knowledge of biology of microorganisms and the microbial engineering techniques need to improve in order to ensure that there are not side effects*.*”* - 2nd hand experience of cancer (18-39 age group).
>
> *“*…*would want certainty that this would not adversely impact anyone else eg future generations*.*”* - 1st hand experience of cancer (60+ age group).

### 2.5 Attitudes towards bacterial therapies

Following the positive overall attitude towards live bacterial therapies, we investigated whether these perceptions changed based on different situations. To this end, we asked participants to rank how comfortable they would be with using a number of different technologies (Figure 4). Most respondents stated they would be comfortable using microbiome engineering technologies in general. However, the majority of respondents stated they would be uncomfortable taking GM foods as part of their diet. This matches with previous responses and studies, highlighting the negative connotations associated with GM foods. The majority of participants stated they would be comfortable using engineered human cells to treat cancer (for example CAR-T therapy) and a similar response was seen towards using ‘engineered’ bacteria to treat cancer. An interesting comment from one participant highlighted how they had not perceived CAR-T cell therapy as a form of genetic engineering at all, potentially leading to them having fewer reservations regarding this kind of treatment over the proposed microbiome engineering techniques presented in the survey. This is perhaps a reminder that the terms used to describe these future treatments greatly impact whether or not they are positively perceived.

**Figure 4.**
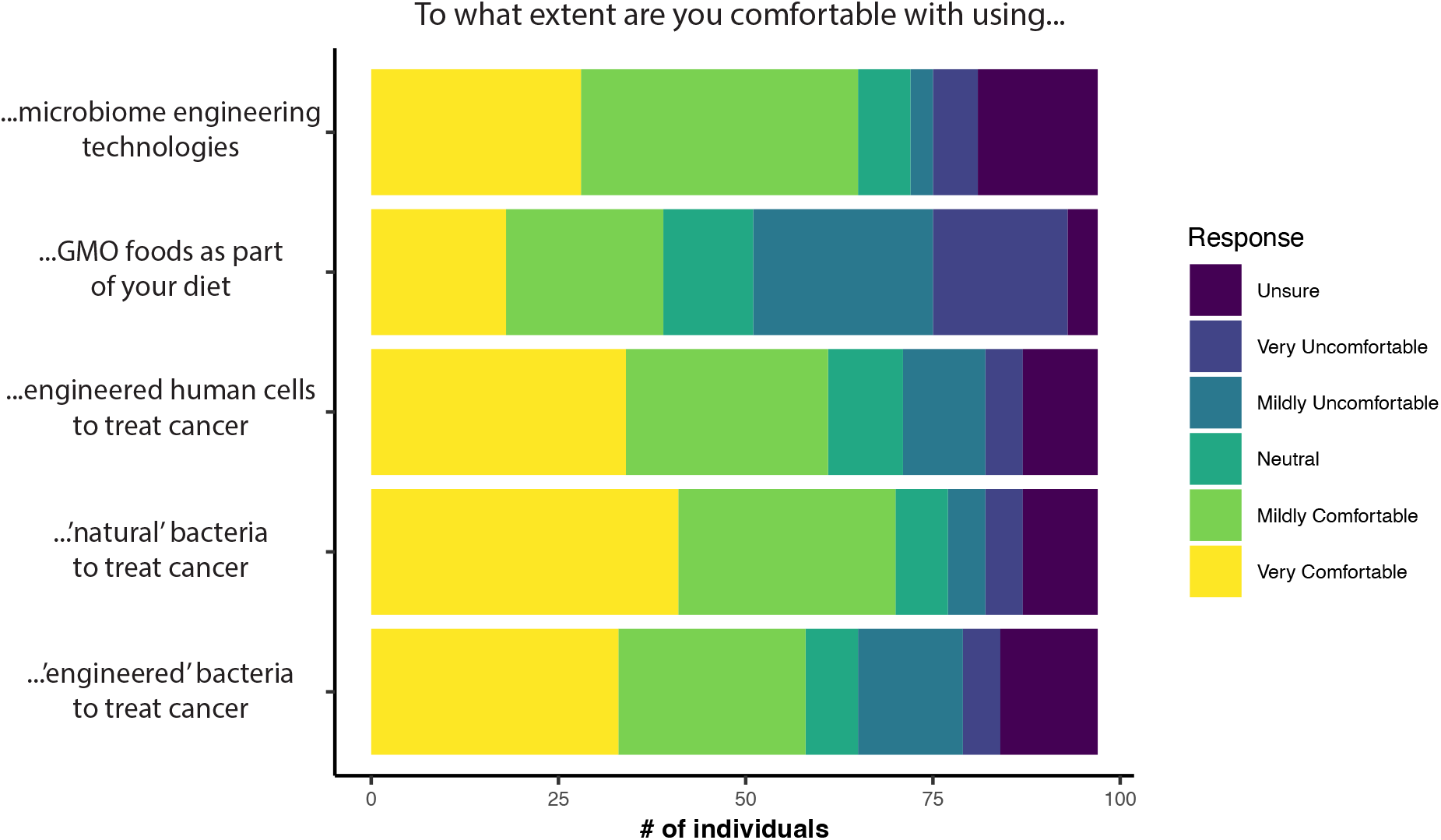
Participants response to the given topics, regarding whether they are comfortable using the highlighted technologies. Answers are coloured by the response given (n = 97 responses).

> *“[I] am very comfy with the concept of CAR-T, funnily enough have never thought of it as genetically modified cells”* - 1st hand experience of cancer (40-59 age group)

Notably, more participants indicated they would be comfortable using ‘natural’ bacterial therapies than either engineering human cells or bacteria. This coincides with previous work by the Nuffield Council on Bioethics that found some people do view natural medicines as safer, healthier and more likely to do good than alternatives (*19*).

Participants who stated they were comfortable with the use of GM foods were found to be more likely to be comfortable with the use of engineered bacteria. There was a positive correlation between those comfortable using engineered human cells and engineered bacteria. No significant correlation was found between age or relationship to cancer and comfort with using engineered bacteria (Table 1). A full breakdown of the statistical analysis performed is given in SI Table S1).

**Table 1.** Positive correlations exist between participants’ opinions on GM foods or engineered human cells and whether they are comfortable with using engineered bacteria to treat cancer. The provided values are p-values from a two-sided T-test, with a p-value below 0.05 indicating statistical significance.

| Are you comfortable using engineered bacteria to treat cancer? |  |
| --- | --- |
| Characteristic | P-value: |
| Age | 0.4946 |
| Relationship with cancer | 1 |
| Comfortable with GM food | 0.0039 |
| Comfortable with engineered human cells | <0.0001 |

Another theme amongst many of the comments was the need for more information. The participants showed a desire to better understand this possible new treatment option. This theme of understanding, whereby participants expressed the desire to know more information regarding this treatment, or that they did not have the knowledge to assess this potential treatment for themselves, was prevalent across the age groups. There is a need from these participants to better understand what these treatments are and a need to correct misinformation. In some responses, there was confusion between GM food and live engineered bacteria. This only serves to highlight how any new treatment needs to be accompanied by information campaigns, and that this is something patients want.

> *“It sounds promising, but i would want to know more about a treatment before accepting it*.*”* - 1st hand experience of cancer (60+ age group)
>
> *“Agree with the theoretical principle but would need to have a lot more info before deciding to use it personally”* - 1st hand experience of cancer (40-59 age group)

The final theme identified centred on optimism towards these new therapies, with 22 comments connected to this theme. Many comments mentioned excitement and hope that these new therapies could bring about improvements in the current standard of cancer care.

> *“Microbiome engineering sounds very promising and far less invasive than some other treatments*.*”* - 2nd hand experience of cancer (60+ age group)
>
> *“I think this would be a huge step forward. Being able to manipulate bacteria in a way which gets the body itself to fight the disease would be incredible*.*”* - 2nd hand experience of cancer (40-59 age group)
>
> *“I’m a great believer in the human body’s ability to use its own resources to counteract disease, and this is mobilising them with perhaps a little modification to do just that*.*”* - 1st hand experience of cancer (60+ age group)
>
> *“Modified bacteria have already shown they hold great potential*.*”* - 1st hand experience of cancer (60+ age group)

## 3 Conclusions

Following the increase in clinical trials that are investigating the efficacy of using genetically modified bacteria to treat cancer, we wished to explore public attitudes towards these technologies. It should be noted that this survey only received a total of 102 full responses and therefore, this limited sample size may not be representative of the wider public in general. In addition, this survey did not collect any data on ethnicity, socio-economic background or geographical location. As such, it is not possible to draw detailed conclusions of how these factors impact on respondents’ willingness to accept these new technologies. Future studies will be needed to explore the influence of these factors in greater detail.

However, the responses still provide a valuable insight into public perceptions surrounding these newly emerging technologies. The majority of participants in this study were comfortable with the concept of live bacterial therapies (both ‘natural’ and ‘engineered’) as cancer treatments, despite a lack of previous knowledge of these terms in some cases. In addition, positive correlations indicated that participants who are comfortable using GM foods or engineered human cells to treat cancer are also more comfortable using engineered bacterial therapies.

To further understand the attitudes of the participants towards microbiome engineering and engineered live bacterial therapies we included open text questions. To identify the main themes of these responses and therefore infer the attitudes of the participants we conducted a thematic analysis. We were able to identify key themes that shape the attitudes of the participants of this survey and support the results of the quantitative questions. For example, trust emerged as a main theme, with many respondents indicating they would be comfortable trusting their healthcare professionals if they recommended microbiome engineering technologies. However, it is important to note the survey could have introduced bias by asking participants about trust in a previous question. Despite this, trust could be an important observation to help guide future avenues to enable successful implementation of new technologies. Overall, these findings are promising for the future adoption of the many microbiome engineering technologies that are currently undergoing clinical trials.

## 4 Methods

### 4.1 Survey Design

As no previous surveys of this nature exist, a bespoke survey design was created using Opinio software. The survey questions were designed to ascertain current attitudes of respondents towards using engineered live bacteria in medicine, specifically as part of cancer treatments. The survey contained 21 questions, including quantitative (e.g. multiple choice ranking) and qualitative (e.g. open-text box) question types. The first section of the survey contained questions relating to current cancer treatments, the second section of the survey assessed participants prior knowledge on probiotics and microbiome engineering, the final section of the survey measured participants comfortability with different concepts, for example CAR-T cell therapies. All patients had to be over the age of 18 and informed consent was obtained with every complete response.

### 4.2 Survey Distribution

The survey was promoted on the Cancer Research UK Patient Involvement webpage (https://www.cancerresearchuk.org/get-involved/volunteer/patient-involvement/involvement-opportunities/survey-can-we-engineer-live-bacteria-to-treat-cancer). The survey was also advertised on email newsletters by the Patient Experience Research Centre (https://www.imperial.ac.uk/patient-experience-research-centre/) and Independent Cancer Patients Voice Network (http://www.independentcancerpatientsvoice.org.uk/). The survey was shared and re-tweeted on X (formerly Twitter). The survey closed to responses on the 31st of December 2022, at 23:59.

### 4.3 Data Analysis

The collected data was downloaded using Opinio software as a csv file and quantitative data analysis performed in R (version: 4.1.2). Incomplete submissions and respondents with no personal connection to cancer (either first or second hand) were removed from further analysis.

For qualitative data, in the form of short text answers, thematic analysis was conducted (*20*). This analysis was performed following the 5 steps of thematic analysis as detailed by Braun and Clarke 2006 (*20*): 1, read the data and identify codes, 2, code the data, 3, identify themes, 4, review the themes, 5, define and name the themes (these themes are given in SI Table S2). An inductive coding approach was used for theme identification.

Further statistical analyses were also performed in R (version: 4.1.2). Firstly, the responses to each question were grouped into comfortable (mildly comfortable and very comfortable), uncomfortable (mildly uncomfortable and very uncomfortable) and unsure (do not know and neutral). A Fisher’s Exact test was then performed on the grouped responses for each relevant question. All conditions tested were assessed against whether the respondent was comfortable with using eLBPs to treat cancer in future.

Data visualisation was performed in RStudio (version: 2022.07.2, (*21*)), using the ggplot2 package(*22*) and Adobe Illustrator (version: 28.5).

## Acknowledgement

The authors …

## References

1. O’Toole, P. W., Marchesi, J. R., and Hill, C. (2017) Next-generation probiotics: the spectrum from probiotics to live biotherapeutics. Nature microbiology 2, 1–6.

2. Lu, T. K., Mimee, M., Citorik, R. J., and Pepper, K. The Chemistry of Microbiomes: Proceedings of a Seminar Series; National Academies Press (US), 2017.

3. Ozdemir, T., Fedorec, A. J., Danino, T., and Barnes, C. P. (2018) Synthetic biology and engineered live biotherapeutics: toward increasing system complexity. Cell systems 7, 5–16.

4. Rutter, J. W., Dekker, L., Owen, K. A., and Barnes, C. P. (2022) Microbiome Engineering: Engineered Live Biotherapeutic Products for Treating Human Disease. Frontiers in Bioengineering and Biotechnology 10, 1735.

5. Gurbatri, C. R., Arpaia, N., and Danino, T. (2022) Engineering bacteria as interactive cancer therapies. Science 378, 858–864.

6. Van den Berghe, L., Masschelein, J., and Pinheiro, V. B. (2024) From competition to cure: the development of live biotherapeutic products for anticancer therapy in the iGEM competition. Frontiers in Bioengineering and Biotechnology 12, 1447176.

7. Miles, S., Ueland, Ø., and Frewer, L. J. (2005) Public Attitudes towards Genetically-modified Food. British Food Journal 107, 246–262.

8. Pardo, R., Midden, C., and Miller, J. D. (2002) Attitudes toward Biotechnology in the European Union. Journal of Biotechnology 98, 9–24.

9. Amin, L., Md Jahi, J., and Md Nor, A. R. (2013) Stakeholders’ Attitude to Genetically Modified Foods and Medicine. The Scientific World Journal 2013, e516742.

10. Woźniak-Gientka, E., Tyczewska, A., and Twardowski, T. (2022) Public Opinion on Biotechnology and Genetic Engineering in the European Union: Polish Consumer Study. BioTechnologia 103, 185–201.

11. Hampel, J., Pfenning, U., and Peters, H. P. (2000) Attitudes towards Genetic Engineering. New Genetics and Society 19, 233–249.

12. Anderson, J., Strelkowa, N., Stan, G.-B., Douglas, T., Savulescu, J., Barahona, M., and Papachristodoulou, A. (2012) Engineering and Ethical Perspectives in Synthetic Biology. EMBO Reports 13, 584–590.

13. Marris, C. (2015) The Construction of Imaginaries of the Public as a Threat to Synthetic Biology. Science as Culture 24, 83–98.

14. EC Study on New Genomic Techniques - European Commission. https://food.ec.europa.eu/plants/genetically-modified-organisms/new-techniques-biotechnology/ec-study-new-genomic-techniques_en.

15. McCarthy, A., Holland, C., and Shapira, P. The Development and Testing of an Early, Rapid Sustainability Assessment Tool for Responsible Innovation in Engineering Biology.

16. Probiotics Industry 2024. https://www.reportlinker.com/market-report/Nutraceutical/11269/Probiotics.

17. Hansen, C. Consumer Insights Findings from New Study of Consumer Understanding of Probiotics and Takeaways for the Food Industry.

18. Lord, K., Ibrahim, K., Kumar, S., Rudd, N., Mitchell, A. J., and Symonds, P. (2012) Measuring Trust in Healthcare Professionals—A Study of Ethnically Diverse UK Cancer Patients. Clinical Oncology 24, 13–21.

19. Nuffield Council on Bioethics, Ideas about Naturalness in Public and Political Debates about Science, Technology and Medicine. 2015.

20. Braun, V., and Clarke, V. (2006) Using Thematic Analysis in Psychology. Qualitative Research in Psychology 3, 77–101.

21. Team, R. C. R: The R Project for Statistical Computing. 2022.

22. Wickham, H. Create Elegant Data Visualisations Using the Grammar of Graphics • Ggplot2. 2022.

