## Supplementary Information for "Engaging cancer patients on their attitudes towards microbiome engineering technologies"

### Supplementary Information:
Engaging cancer patients and their families on aspects of microbiome engineering

**3 Survey Questions** **4**

### 1 Extended materials & methods

#### 1.1 Survey definitions

Definitions of key concepts explored within the survey were provided for all participants. These definitions are provided below.

**Bacteria:** Bacteria are small organisms, or microscopic living things, that can be found in nearly all natural environments. They live on our skin and inside our bodies. Some bacteria may cause infection and disease. However, many other bacteria help us to digest our food, produce vitamins (such as B12 and K) and fight off other bacteria.

**Probiotics:** Probiotics are live microorganisms (such as bacteria) that are intended to have health benefits when consumed or applied to the body. They can be found in fermented foods, dietary supplements, and beauty products. Probiotics have so far shown some promise in the treatment of diarrhoea, bacterial vaginosis, and irritable bowel syndrome (IBS).

**Microbiome engineering:** A microbiome is a community of microorganisms (bacteria, fungi, etc.) that live together within a specific environment, like the human gut. Microbiome engineering aims to modify these communities in a predictable way, for example, for the treatment of diseases within the body. There are currently many ongoing clinical trials for the treatment of disease with live microorganisms.

**GMO:** A genetically modified organism (GMO) is any organism which has had its genetic material, or DNA, modified in a laboratory. Genetically modified crops are routinely produced and consumed in the USA, though not yet in the UK and Europe. As microbiome engineering may involve the use of GMOs, the following questions will explore your opinions on the use of GMOs.

### 2 Extended results

#### 2.1 Survey participant demographics

A full breakdown of the participant demographics is given in Figure S1. As noted in the main text, the responses of participants with 'no personal' relationship to cancer were excluded from further analysis.

#### 2.2 Statistical analysis of quantitative data

Statistical analysis was performed to identify positive correlations between the answers received across different survey questions. This analysis identified positive correlations between participants who identified as comfortable with GM foods/engineered human cells and those who were comfortable with using engineered bacteria to treat cancer (SI Table 1).

Table S1: Identification of positive correlations between survey answers and whether the participant is comfortable using engineered bacteria to treat cancer (a p-value < 0.05 was considered statistically significant).

| | **Are you comfortable using engineered bacteria?** | | |
|---|---|---|---|
| | Number of participants (%) | | |
| **Characteristic** | **Comfortable (n=58)** | **Uncomfortable (n=19)** | **p-value** |
| **Age** | | | 0.4946 |
| 18 – 39 | 12 (%) | 3 (%) | |
| 40 – 59 | 15 (%) | 5 (%) | |
| 60+ | 31 (%) | 11 (%) | |
| Fisher's Exact Test, Alt. hypothesis: 2 sided | | | |
| **Relationship with cancer** | | | 1 |
| First-hand relationship | 33 (75%) | 11 (25%) | |
| Second-hand relationship | 25 (76%) | 8 (24%) | |
| Fisher's Exact Test, Alt. hypothesis: 2 sided | | | |
| **Comfortable with GMO** | | | 0.0039 |
| Comfortable | 30 (94%) | 2 (6%) | |
| Neutral | 2 (100%) | 0 (0%) | |
| Uncomfortable | 26 (60%) | 17 (40%) | |
| Fisher's Exact Test, Alt. hypothesis: 2 sided | | | |
| **Comfortable with engineered human cells** | | | <0.0001 |
| Comfortable | 51 (93%) | 3 (7%) | |
| Neutral | 7 (78%) | 2 (22%) | |
| Uncomfortable | 0 (0%) | 14 (100%) | |
| Fisher's Exact Test, Alt. hypothesis: 2 sided | | | |

##### 2.3 Themes from thematic analysis

| Theme | Definition |
|---|---|
| Trust | Explicit mentions of trusting health care professionals and their recommendations. |
| Social connection | Considering the impact of this technology on the self and on other people. |
| Optimism | Positive tone in responses towards microbiome engineering. There is excitement and hope towards these technologies. |
| Understanding | Participants express their lack of knowledge or desire for more information on a topic that is new to them. |

Table S2: Themes and their definitions identified during the thematic analysis of the qualitative open text responses in the survey.

#### 3 Survey Questions

##### Section 1

**Q1.** Please select your age range from the groups below:

- A. 18-39
- B. 40-59
- C. 60+
- D. Prefer not to say

**Q2.** How have you personally been impacted by cancer?

- A. 1st hand (i.e., current patient/survivor of cancer)
- B. 2nd hand (i.e., first-degree relative of a patient who has been diagnosed with cancer)
- C. 3rd hand (i.e., friend/distant relative of a patient who has been diagnosed with cancer)
- D. Not personally impacted

**Q3.** Which of these potential cancer treatment options would you be comfortable taking if recommended (or have previously taken if applicable)?

- A. Surgery – removal of tumours through cutting them out of the body.
- B. Chemotherapy – use of drugs to kill cancer cells.
- C. Radiation – use of high-powered energy rays (such as X-rays) to kill cancer cells.
- D. Immunotherapy/Biological Therapy – use of the body's own immune system to fight cancer cells.

- E. Live bacterial therapy – use of live bacteria to target cancer inside the body.
- F. Do not know enough to answer.

**Q4.** Who would you trust to provide information on new cancer therapies and recommendations for promising treatments? Please rank these options from most trustworthy (1) to least trustworthy (6):

- A. Clinicians/doctors
- B. Regulatory bodies (e.g., Medicines & Healthcare products Regulatory Agency, European Medical Agency)
- C. Cancer charities
- D. NHS website and official resources (e.g., hospital flyers and brochures)
- E. Academic researchers
- F. Pharmaceutical/private research companies

#### Section 2

You will now be given a short description of the term 'Probiotics'. Please read the following statement carefully:

Probiotics are live microorganisms (such as bacteria) that are intended to have health benefits when consumed or applied to the body. They can be found in yogurt and other fermented foods, dietary supplements, and beauty products. Probiotics have so far shown some promise in the treatment of diarrhoea, bacterial vaginosis, and irritable bowel syndrome (IBS).

**Q5.** Please select the statement that best applies to you regarding probiotics:

- A. I had never heard of probiotics and I do not understand what they are.
- B. I had never heard of probiotics but I understand what they are now.
- C. I have heard about probiotics previously but do not understand what they are.
- D. I have heard about probiotics previously and I understand what they are.

**Q6.** Are you aware of the proposed benefits of probiotics?

- A. Yes
- B. No
- C. Do not know enough information to provide an opinion.

**Q7.** Are you aware of the limitations of probiotics?

- A. Yes

- B. No
- C. Do not know enough information to provide an opinion.

**Q8.** Would you/do you take probiotics to supplement your current diet? This includes regularly consuming items such as fermented foods or live yogurt.

- A. Yes
- B. No
- C. Do not know enough information to provide an opinion.

**Q9.** Would you/do you take probiotics to treat a disease/medical condition?

- A. Yes
- B. No
- C. Do not know enough information to provide an opinion.

#### Section 3

You will now be given a short description of the term 'microbiome engineering'. Please read the following statement carefully:

A microbiome is a community of micro-organisms (bacteria, fungi, etc.) that live together within a specific environment, like the human gut. Microbiome engineering aims to modify these communities in a predictable way, for example, for the treatment of diseases within the body.

**Q10.** Please select the statement that best applies to you regarding microbiome engineering:

- A. I had never heard of it and I do not understand what it is.
- B. I had never heard of it but I understand what it is now.
- C. I have heard about it previously but do not understand what it is.
- D. I have heard about it previously and I understand what it is.

**Q11.** Please rate to what extent you would approve/disapprove of microbiome engineering technologies:

- A. Strongly disapprove
- B. Mildly disapprove
- C. Neutral/no opinion
- D. Mildly approve

- E. Strongly approve
- F. Do not know enough to answer

Please explain your choice in a maximum of 100 words (this box can be left blank, please DO NOT include any information that could potentially identify you, i.e., name, exact age, where you live, health status, etc.):

*[Open text box for responses]*

**Q12.** If you disapprove of microbiome engineering, please highlight your major concerns and what, if anything, could be done to address them? (Please DO NOT include any information that could potentially identify you, i.e., name, exact age, where you live, health status, etc.):

*[Open text box for responses]*

You will now be given a short description of the term ‘genetically modified organism’. Please read the following statement carefully:

A genetically modified organism (GMO) is any organism which has had its genetic material, or DNA, modified in a laboratory. As microbiome engineering may involve the use of GMOs, the following questions will explore your opinions on the use of GMOs.

**Q13.** Please rate to what extent you approve/disapprove of consuming genetically modified foods (e.g., GM tomatoes) as part of your diet:

- A. Strongly disapprove
- B. Mildly disapprove
- C. Neutral/no opinion
- D. Mildly approve
- E. Strongly approve
- F. Do not know enough to answer

**Q14.** Please rate how comfortable you would be with using engineered (i.e., genetically modified) human cells as part of your cancer treatment plan (e.g., currently available CAR-T cell therapy):

- A. Very uncomfortable
- B. Mildly uncomfortable
- C. Neutral/no opinion
- D. Mildly comfortable
- E. Very comfortable
- F. Do not know enough to answer

**Q15.** In the future, how comfortable would you be with using natural live bacteria as part of your cancer treatment plan:

- A. Very uncomfortable
- B. Mildly uncomfortable
- C. Neutral/no opinion
- D. Mildly comfortable
- E. Very comfortable
- F. Do not know enough to answer

**Q16.** In the future, how comfortable would you be with using engineered (i.e., genetically modified) live bacteria as part of your cancer treatment plan:

- A. Very uncomfortable
- B. Mildly uncomfortable
- C. Neutral/no opinion
- D. Mildly comfortable
- E. Very comfortable
- F. Do not know enough to answer

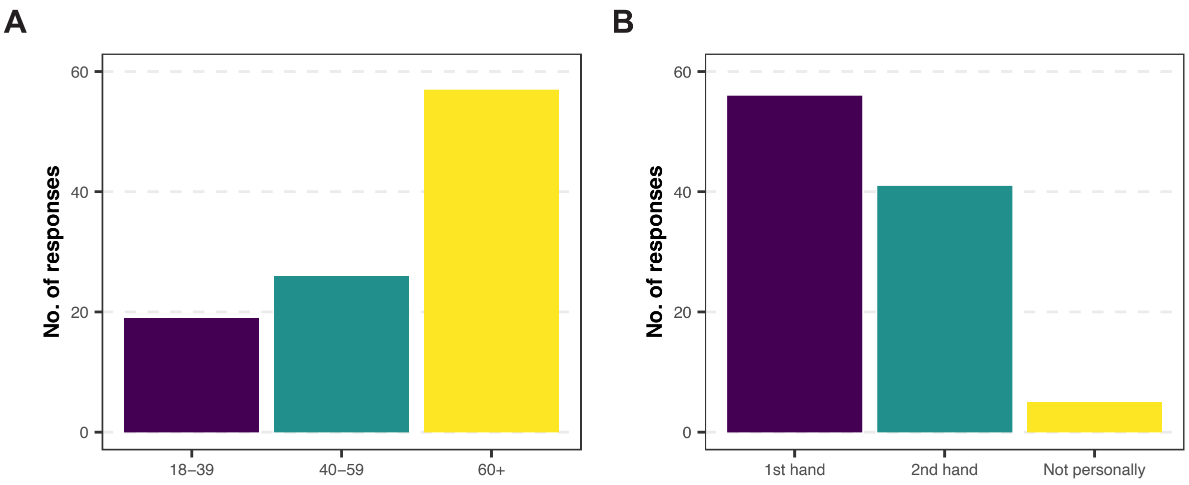

Figure S1: Participant demographics of the 102 total responses. (**A**) Age group of participant, (**B**) Participant relationship with cancer ($1^{st}$ hand = patient/survivor, $2^{nd}$ hand = family/friend of a cancer patient).
